# Biogeographic patterns of ectomycorrhizal fungal communities associated with *Castanopsis sieboldii* across the Japanese archipelago

**DOI:** 10.1101/632901

**Authors:** Shunsuke Matsuoka, Takaya Iwasaki, Yoriko Sugiyama, Eri Kawaguchi, Hideyuki Doi, Takashi Osono

**Author notes:** Author for correspondence: Shunsuke Matsuoka.

## Abstract

Biogeographic patterns in ectomycorrhizal (ECM) fungal communities and their drivers have been elucidated, including effects of host tree species and abiotic (climatic and edaphic) conditions. At these geographic scales, genotypic diversity and composition of single host tree species change with spatial and environmental gradients, reflecting their historical dispersal events. However, whether the host genotypes can be associated with the biogeographic patterns of ECM communities remains unclear. We investigated the biogeographic pattern of ECM fungal community associated with the single host species *Castanopsis sieboldii* (Fagaceae), whose genotypic diversity and composition across the Japanese archipelago has already been evaluated, and we quantified the effect of host genotypes on the biogeographic pattern. Richness and community composition of ECM fungi changed with latitude and longitude; these biogeographic changes of ECM community were significantly explained by host genotypic variables. Quantitative analyses showed a higher relative explanatory power of climatic and spatial variables than that of host genotypic variables for the biogeographic patterns in the ECM community. Our results suggest the importance of historical events of host dispersal in determining the biogeographic patterns of the ECM fungal community, while their explanation power was lower than that for climatic filtering and/or fungal dispersal.

## Introduction

Ectomycorrhizal (ECM) fungi are symbionts of tree species in Fagaceae, Betulaceae, Dipterocarpaceae, and Pinaceae and are dominant in tropical, temperate, and boreal forests (Brundrett, 2009). The ECM fungi are a major component of the forest floor and play an essential role in nutrient cycling via exchanging soil nutrients and therefore are critical for determining and maintaining forest ecosystem processes (Smith & Read, 2008). To infer the community responses to environmental changes, the relationships between spatial patterns of ECM fungal communities and factors responsible for those patterns have been investigated in previous studies (Lilleskov & Parrent, 2007). In these studies the spatial variations of ECM fungal community were related to key environmental factors including both biotic, such as species identity and phylogeny of the host (Ishida *et al.*, 2007; Tedersoo *et al.*, 2013), and abiotic factors, such as soil property (like soil pH) and climatic conditions (e.g., Bahram *et al.*, 2012; Horton *et al.*, 2013). Whereas, recent studies have shown that dispersal of ECM fungi is limited spatially, even at a few kilometers, and that this dispersal limitation can generate spatial structures in ECM fungal communities independent of environmental factors (Peay *et al.*, 2012; Peay & Bruns, 2014). Thus far, the effects of these factors have been investigated at relatively small spatial scales, from forest to landscape scales (e.g., Tedersooet *et al.*, 2011; Bahram *et al.*, 2012; Matsuoka *et al.*, 2016a).

Recently, in some pioneer studies the geographic distributions of ECM fungal communities at regional and/or continental scales were investigated (ranging from hundreds to thousands of kilometers). From these studies, it was revealed that species identity or phylogeny of host trees are one of the most important factors generating geographic structures of ECM fungal communities (Põlme *et al.*, 2013; Roy et al. 2013; van der Linde *et al.*, 2018; Wu *et al.*, 2018). Unlike free-living organisms, host-associated microbial communities like ECM fungi often exhibit biogeographic patterns that are related to the distribution of their hosts (Martiny *et al.*, 2006), because of co-migration of microbes with their host species to new habitats (Kennedy *et al.*, 2011). Therefore, these recent results imply that some part of ECM fungal communities respond to environmental change by following the movement of host trees. However, in studies at regional and/or continental scales, host tree compositions inevitably show the biogeographical structure reflecting both present environments and historical distribution and dispersal (e.g., Svenning & Skov 2007; Jiménez-Alfaro *et al.*, 2018). Therefore, in studies of large spatial scales, within which host species composition changes with geographical gradients, whether the observed biogeographic patterns of ECM fungal communities are related to the host biogeography or whether the ECM fungal community itself is responding to other factors, such as environmental and spatial factors, has not been always distinguished appropriately (Miyamoto *et al.*, 2015; Matsuoka *et al.*, 2016a). Therefore, in order to evaluate how much of the geographic pattern of ECM fungal communities are attributable to the fungal response to environment or dispersal limitation, studies focusing on single host species are indispensable.

When focusing on a single host species, especially those that are distributed across climatic zones, it is important to note that the genotypic structures of host species between geographically proximal sites can resemble each other. Distribution ranges of most tree species have expanded and moved from the refugia in the Last Glacial Maximum. Thus, genotypes of individual host species have their own geographic structure (e.g., genotypic composition and diversity), called phylogeography. Such phylogeographic patterns of host species (e.g., differences in genotypes across site) can also influence the ECM fungal communities, as some evidence show that ECM fungi co-migrate with their hosts (Kennedy *et al.*, 2011) or they show preference to certain intra-specific genotypes (Gehring *et al.*, 2017; Patterson *et al.*, 2019). Thus, when the distribution patterns of ECM fungal communities correspond with host genetic pattern, there are two possible explanations for the correspondence: (1) Such patterns were generated in relation with their host species (e.g., co-migration) and (2) they were generated owing to the effects of direct fungal response to environment and dispersal limitation, independent to the host. However, so far, the relationships between geographic patterns of host genotypic structures and geographic distribution of ECM communities, and quantitative evaluation of the relative importance of the two possible explanations for the distribution pattern of ECM communities have been investigated in few studies.

We aimed to evaluate the geographic pattern of ECM fungal community and the effect of host genotypes and genotypic diversity on that pattern by focusing on a single host species *Castanopsis sieboldii* (Makino) Hatus. ex T.Yamaz. et Mashiba (Fagaceae). We chose *C. sieboldii* as the focal host species because this species is widely distributed across different climate regions of the Japanese archipelago and its phylogeographic pattern over its distribution range has been already documented with genetic markers (Aoki *et al.*, 2014). Importantly, unlike Europe and North America, ice sheets were not present in the Japanese archipelago during the Quaternary period, and this, together with the complex mountainous terrain of Japan, resulted in the establishment of tree refugia in multiple regions across Japan (e.g. Tsukada, 1984). Therefore, the present distributions of individual genotypes of single species are assumed to reflect the environmental gradients like climate (Tsukada, 1984; Aoki *et al.*, 2014). For these features, we considered *C. sieboldii* in Japan as the appropriate host species to evaluate the effect of host genotypes and genotypic diversity on the geographic distribution of the associated ECM fungal community. We specifically hypothesized that the richness and composition of ECM fungal communities change with geographic gradient (i.e., latitude and longitude) and that the geographic variation is affected by host genetic variables. We further quantified the relative effects of environmental factors (i.e., soil property and climate) and spatial factors on the geographic pattern of ECM fungal community to identify the effects of host genetic variables.

## Materials and Methods

### Study sites and sampling procedure

We focused on an ECM host tree species *Castanopsis sieboldii* (Fagaceae), a dominant, canopy, and climax species in the Japanese *Castanopsis*-type evergreen broad-leaved forests. These *Castanopsis*-type forests characterize the biodiversity and endemism of the subtropical and warm-temperate zone in Japan. Sampling was conducted in 12 mature *Castanopsis*-dominated forests (Table 1 and Fig. 1) covering almost the entire distribution range of *C. sieboldii*, from Site 1 in Ryukyu Islands (~25°N) adjacent to the present southern boundary of the host species’ distributional range to Site 12 in Sado Island (~38°N) adjacent to its northern boundary. Latitudes and longitudes of the sampling sites were strongly correlated (Pearson’s *r* = 0.845, P < 0.001), as the Japanese Archipelago is elongated from northeast to southwest. Sampling was conducted once for each site in the summer (from July to early September) in 2011, 2012, and 2013. At each site, sampling was conducted in < 1 ha stands dominated by *Castanopsis sieboldii* (Fagaceae). We selected 20 individuals of *C. sieboldii* (diameter at breast height > 20 cm) and collected a block of fermentation-humus (FH) layer (10 × 10 cm, 10 cm in depth) including tree roots within 3 m from each tree trunk. The soil blocks were collected within the expanse of the canopies of each individual *C. sieboldii* tree and the sampling points were at least 3 m from each other. The blocks were stored in plastic bags and frozen at −20°C during transport to the laboratory. A total of 240 blocks (12 study sites × samples from 20 individual of *C. sieboldii*) were used for the study.

**Table 1.** The location and environmental factors of the study sites.

| Site No. | Site name | n<sup>1</sup> | Latitude (°N) | Longitude (°E) | sampling | pH<sup>2</sup> | WC<sup>2</sup> | CN<sup>2</sup> | MAT (°C) | MAP (mm) | t2w (°C) | p2w (mm) |
| --- | --- | --- | --- | --- | --- | --- | --- | --- | --- | --- | --- | --- |
| 1 | Ishigaki | 20 | 24.4178 | 124.1881 | Jul. 2013 | 4.10 ± 0.36 | 29.7 ± 5.6 | 17.8 ± 2.0 | 24.4 | 2130.0 | 29.8 | 217.0 |
| 2 | Amami | 20 | 28.2225 | 129.3412 | Jul. 2013 | 3.24 ± 0.31 | 30.9 ± 14.6 | 21.8 ± 4.0 | 21.9 | 2420.5 | 28.6 | 0.0 |
| 3 | Yakushima | 20 | 30.2575 | 130.5817 | Jul. 2013 | 3.45 ± 0.41 | 65.1 ± 13.6 | 21.3 ± 2.5 | 20.4 | 3186.8 | 26.5 | 239.5 |
| 4 | Hitoyoshi | 20 | 32.1491 | 130.7803 | Jul. 2012 | 4.16 ± 0.14 | 48.6 ± 10.5 | 14.0 ± 1.7 | 15.8 | 2535.3 | 24.7 | 460.0 |
| 5 | Saeki | 20 | 32.9594 | 131.8913 | Jul. 2012 | 3.52 ± 0.57 | 56.8 ± 12.4 | 17.1 ± 1.8 | 16.8 | 2260.8 | 25.1 | 216.5 |
| 6 | Ashizuri | 16 | 32.7447 | 133.0002 | Jul. 2012 | 3.88 ± 0.28 | 59.5 ± 13.9 | 16.0 ± 2.2 | 18.4 | 2585.3 | 26.1 | 75.0 |
| 7 | Kii | 19 | 33.4653 | 135.8333 | Aug. 2011 | 3.57 ± 0.26 | 41.0 ± 11.5 | 15.6 ± 1.4 | 17.4 | 2700.0 | 27.0 | 35.0 |
| 8 | Hachijo | 20 | 33.1076 | 139.8414 | Aug. 2012 | 4.20 ± 0.45 | 49.7 ± 8.7 | 15.8 ± 1.0 | 18.0 | 3324.8 | 26.5 | 9.5 |
| 9 | Izu | 20 | 34.6509 | 138.8526 | Aug. 2011 | 3.61 ± 0.22 | 34.3 ± 8.5 | 16.7 ± 1.1 | 16.8 | 1748.6 | 25.6 | 51.5 |
| 10 | Chiba | 18 | 35.0934 | 139.9162 | Aug. 2011 | 4.02 ± 0.35 | 21.6 ± 7.9 | 14.4 ± 2.2 | 16.1 | 1918.0 | 26.7 | 9.0 |
| 11 | Kurayoshi | 20 | 35.4248 | 133.8224 | Sep. 2012 | 3.61 ± 0.22 | 35.2 ± 13.7 | 18.5 ± 2.4 | 14.9 | 1791.2 | 27.4 | 23.0 |
| 12 | Sado | 20 | 37.9661 | 138.3680 | Sep. 2013 | 3.78 ± 0.27 | 41.6 ± 8.4 | 18.2 ± 2.7 | 13.8 | 1846.1 | 25.5 | 146.0 |

<sup>1</sup> number of sample used for analyses (see text), <sup>2</sup> Mean ± S.D. pH pH of FH (fermentation-humus) materials, WC water content of FH materials, CN C/N ratio of FH materials, MAT mean annual temperature, MAP mean annual precipitation, t2w mean daily temperature during the 2 weeks before each sampling date, p2w cumulative amount of precipitation during 2 weeks before each sampling date.

**Fig. 1.**
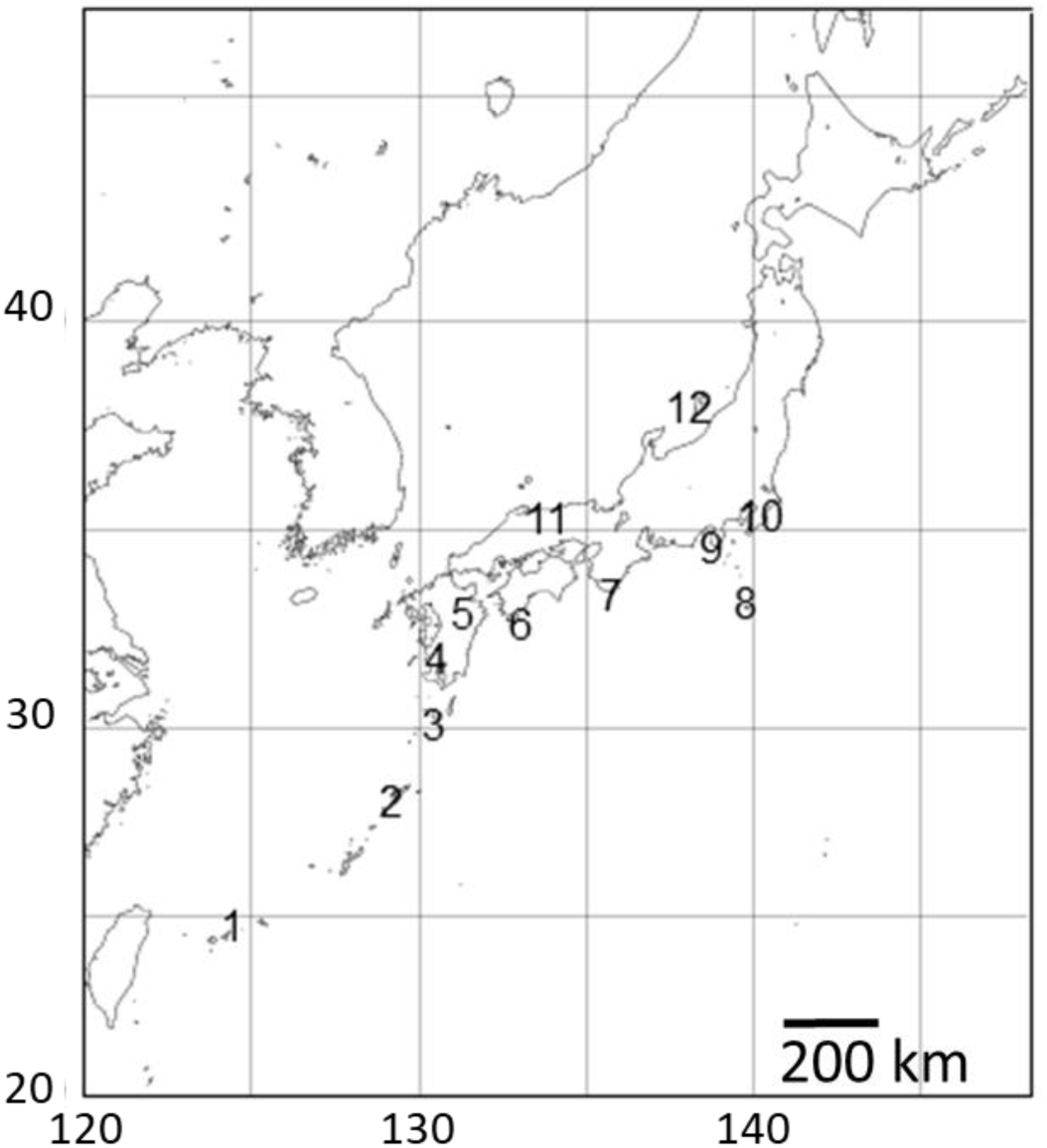
Sampling sites in this study. Numbers are consistent with site number listed in Table 1. Axes indicate latitude °N (y) and longitude °E (x).

In the laboratory, FH materials in the blocks were slightly melted at room temperature (20-25°C) and gently loosened. Then fine roots of trees were extracted from the samples using a 2-mm mesh sieve and gently washed with tap water to remove soil particles and debris. The sieved FH materials were used for the measurement of chemical properties as described below. In each block, 20 individual root segments (approximately 5 cm in length) were selected, and one root tip (1 to 2 mm in length) was collected from each root segment under a binocular microscope at 20× magnification. The 20 root tips were pooled for each block and kept in a microcentrifuge tube containing 70% ethanol (w/v) at −20°C. Before extracting DNA, the root tips were washed to remove small particles on the root surface by 0.005% aerosol OT (di-2-ethylhexyl sodium sulfosuccinate) solution (w/v) and rinsed with sterile distilled water. The root tips were then transferred to tubes containing cetyltrimethylammonium bromide (CTAB) lysis buffer and stored at −20°C until DNA extraction.

### DNA extraction, PCR amplification, and pyrosequencing

The methods of DNA analysis were generally according to those described in Matsuoka *et al.* (2016b). Whole DNA was extracted from root tips in 240 samples using the modified CTAB method described by Gardes & Brum (1993). For direct 454 sequencing of the fungal internal transcribed spacer 1 (ITS1), we used a semi-nested PCR protocol. The ITS region has been proposed as the standard fungal barcode (Schoch *et al*., 2012). First, the entire ITS region and the 5′-end region of large sub unit (LSU) were amplified using the fungus-specific primers ITS1F (Gardes & Bruns, 1993) and LR3 (Vilgalys & Hester, 1990). Polymerase chain reaction was performed in a 20 μl volume with the buffer system of KOD FX NEO (TOYOBO, Osaka, Japan), which contained 1.6 μl of template DNA, 0.3 μl of KOD FX NEO, 9.0 μl of 2× buffer, 4.0 μl of dNTP, 0.5 μl each of the two primers (10 μM) and 4.1 μl of distilled water. The PCR conditions were as follows: an initial step of 5 min at 94°C; followed by 23 cycles of 30 s at 95°C, 30 s at 58°C for annealing, and 90 s at 72°C; and a final extension of 10 min at 72°C. The PCR products were purified using ExoSAP-IT (GE Healthcare, Little Chalfont, Buckinghamshire, U.K.) and diluted by adding 225 μl of sterilized water. The second PCR was then conducted targeting the ITS1 region using the ITS1F fused with the 454 Adaptor and the eight base-pair DNA tag (Hamady *et al*., 2008) for post-sequencing sample identification and the reverse universal primer ITS2 (White *et al.*, 1990) fused with the 454 Adaptor. The PCR was performed in a 20 μl volume with the buffer system of KOD Plus NEO (TOYOBO), which contained 1.0 μl of template DNA, 0.2 μl of KOD Plus NEO, 2.0 μl of 10× buffer, 2.0 μl of dNTP, 0.8 μl each of the two primers (5 μM), and 13.2 μl of distilled water. The PCR conditions were as follows: an initial step of 5 min at 94°C; followed by 28 cycles of 30 s at 95°C, 30 s at 60°C, and 90 s at 72°C; and a final extension of 10 min at 72°C. The PCR products were purified with ExoSAP-IT and quantified with Nanodrop. Amplicons were pooled into five libraries and purified using an AMPure magnetic bead kit (Beckman Coulter, California, USA). The pooled products were sequenced in five 1/16 GS-FLX sequencer (Roche 454 Titanium) at the Graduate School of Science, Kyoto University, Japan.

### Bioinformatics analyses

The procedures used for bioinformatics analyses followed those described in Matsuoka *et al.* (2016a). In pyrosequencing, 315,685 reads were obtained. These reads were trimmed with sequence quality (Kunin *et al.*, 2010) and sorted into individual samples using the sample-specific tags. The remaining 205,261 reads were deposited in the Sequence Read Archive of DNA Data Bank of Japan (accession: DRA004281). The pyrosequencing reads were assembled using Claident pipeline v0.1.2013.08.10 (Tanabe & Toju, 2013; software available online: https://www.claident.org/), which is a highly parallelized extension of the Minimus assembly pipeline (Sommer *et al.*, 2007). We removed the short reads (<150 bp) and then removed potentially chimeric sequences and pyrosequencing errors using UCHIME v4.2.40 (Edger *et al.*, 2011) and algorithm in CD-HIT-OTU (Li *et al.*, 2012), respectively. After these filtering procedures, 130,004 reads were obtained. Reads from seven samples that had less than 50 filtered reads, were not used in the following analyses because such low read numbers could lead to the underestimation of ITS richness. Thus, the remaining 129,725 reads from 233 samples were used for further analyses. The number of sequencing reads per sample ranged from 50 to 3058 (mean, 551). All sequences were assembled across the samples using Assams at a threshold similarity of 97%, which is widely used for fungal ITS region (Osono, 2014), and the resulting consensus sequences represented molecular operational taxonomic units (OTUs). Then singleton OTUs were removed. Consensus sequences of the OTUs are listed in (Supporting Information Table S1).

To systematically annotate the taxonomy of the OTUs, we used Claident v0.1.2013.08.10 (Tanabe & Toju, 2013), built upon automated BLAST-search by means of BLAST+ (Camacho *et al.*, 2009) and the NCBI taxonomy-based sequence identification engine. Using the reference database from INSDC for taxonomic assignment, sequences homologous to ITS sequence of each query were fetched, and then taxonomic assignment was performed based on the lowest common ancestor algorithm (Huson *et al.*, 2007). The results of Claident and the number of reads for the OTUs are given in Table S1. To screen for ECM fungi, we referred to reviews by Tedersoo & Smith (2013) to assign OTUs to the genera and/or families that were predominantly ECM fungi. The resultant ECM fungal OTUs (ECM OTUs) were used for further analyses (see Supporting Information Table S1).

### Environmental and host genetic data

To investigate relationships between environmental factors (i.e., edaphic, climatic, and host biogeographic variables) and ECM fungal community (richness and composition of ECM OTUs), we measured and calculated the following variables (Table 1 and Supporting Information Method S1). For each sample, water content (WC), pH, and carbon (C) and nitrogen (N) concentrations of the FH materials were measured. Carbon and nitrogen concentrations were determined by the combustion method using automatic gas chromatography (Sumigraph NC-22, Sumika Chemical Analysis Service, Ltd, Tokyo, Japan). The methods for WC, pH, and C and N concentrations followed those described in Matsuoka *et al.* (2016a). We calculated four climatic variables for each site: the mean annual temperature (MAT) (10-year averages before each sampling date), the mean annual precipitation (MAP) (10-year averages before each sampling date), the mean daily temperature during the 2 weeks before each sampling date (t_2w_), and the cumulative amount of precipitation during the 2 weeks before each sampling date (p_2w_). Data of air temperature and rainfall were obtained from the nearest station of the Automatic Metrological Data Acquisition System (Japan Meteorological Agency) to each study site. All stations are located within 8 km from each study site.

For evaluation of effects of the host genotypes and genotypic diversity on the ECM community, we used three host genotype variables and three host genetic diversity variables that were estimated from the data published by Aoki *et al.* (2014). Aoki *et al.* (2014) conducted phylogeographic analysis with 32 microsatellite primers (expressed sequence tags-simple sequence repeats, EST-SSRs), targeting 63 populations of *C. sieboldii* in Japan. Membership probabilities of three STRUCTURE clusters were used as host genotype variables. STRUCTURE analysis is a model-based Bayesian clustering approach to estimate genetic differentiation based on multi-locus genotype data (Pritchard *et al.*, 2000; Falush *et al.*, 2003). This method has been widely used for many population genetic and phylogeographic studies. Aoki *et al.* (2014) have reported that three genetic clusters were clearly detected within *C. sieboldii*, which are distributed in Ryukyu, western Japan, and eastern Japan, respectively (see also Supporting Information Method S1). The combination of these host genotype variables can represent the genetic differentiation pattern within the host species at each sampling site. The three host genetic diversity variables were allelic richness, frequencies of rare alleles, and frequency of private alleles. Allelic richness is a parameter which represents genetic diversity (El Mousadik & Petit, 1996). The percentage of rare (less than 5% in total) and private (endemic to one population) alleles represent genetic uniqueness. Rare alleles show relatively high sensitivity and private alleles indicate high uniqueness. Historical stabilized populations such as refugia during glacial periods show high genetic uniqueness, and admixture populations derived from multiple refugia indicate high genetic diversity and low genetic uniqueness (Petit *et al.*, 2003). Therefore, considering these three host genetic diversity variables and three host genotype variables mentioned above, we could evaluate the effect of host genotypes and genotypic diversity. Details of data processing and all host genetic variables are described in Supporting Information Method S1 and Table S2. The correlations between latitude and longitude, and each environmental variable are provided in Supporting Information Table S3.

### Data analyses

The presence or absence of the ECM OTUs was used for all data analyses as binary data regardless of the number of 454 reads, because there are known issues with quantitative use of read numbers generated from amplicon sequencing (Amend *et al.*, 2010; Elbrecht & Leese, 2015). All analyses were performed using R v. 3.0.1 (R Development Core Team, 2013).

Differences in the sequencing depth of individual samples affect the number of OTUs retrieved, often leading to the underestimation of OTU richness in those samples that had low sequence reads. In our dataset, because the rarefaction curves for some samples did not reach asymptotes (Supporting Information Fig. S1), we conducted coverage-based rarefaction (Chao & Jost, 2012) to cover 96% of the total diversity of each sample. These rarefied numbers were used in the analyses of OTU richness. The rarefied OTU numbers were strongly related to the raw OTU numbers per sample (Pearson’s *r* = 0.890, P < 0.001, Supporting Information Table S4).

To examine the biogeographic pattern of the rarefied ECM OTU richness, Pearson’s correlations between geographic variables, or latitude and longitude, and OTU richness were evaluated. Then we analyzed the relationship between the ECM OTU richness and the environmental variables using a generalized linear model (GLM). Error structure and link function of the GLM was Poisson and log, respectively. Environmental variables included 10 variables, that is three edaphic variables (pH, C/N ratio, and WC), four climatic variables (MAT, MAP, t2w, and p2w), and three host genetic diversity variables (allelic richness, frequency of rare alleles, and frequency of private alleles). The best model to explain the variation in OTU richness was chosen using the backward model selection based on Bayesian information criterion (BIC). For all variables in the selected model, partial pseudo-R^2^ were calculated according to Nagelkerke (1991).

To examine the biogeographic pattern of the ECM OTU composition, correlations between geographic variables, or latitude and longitude, and ECM OTUs composition were evaluated using the Mantel test with 9999 permutations. Geographic variables and presence/absence data of ECM OTUs for each sample were converted into a dissimilarity matrix using Euclidian distance and the Raup-Crick dissimilarity index (‘raupcrick’ command in the vegan R package), respectively. The Raup-Crick dissimilarity index is a probabilistic index and is less affected by the species richness gradient among sampling units in comparison to other major dissimilarity indices such as Jaccard’s and Sørensen’s indices (Chase *et al.*, 2011). Then community dissimilarity of ECM OTUs among sites was ordinated using nonmetric multidimensional scaling (NMDS). Presence/absence data of ECM fungal OTUs for each site were merged and converted into a dissimilarity matrix using the Raup-Crick index. Correlation of NMDS structure with geographic coordinates (latitude and longitude) were tested by permutation tests (‘envfit’ command in the vegan package, 9999 permutations).

We used variation partitioning based on the distance-based redundancy analysis (db-RDA, ‘capscale’ command in the vegan package) to quantify the contribution of the edaphic, climatic, host genetic, and spatial variables to the community structure of ECM fungal OTUs. The relative weight of each fraction (purely and shared fractions and unexplained fractions) was estimated following the methodology described by Peres-Neto *et al.* (2006). For the db-RDA, the presence/absence OTUs data for each sample were converted into a Raup-Crick dissimilarity matrix, and we constructed four models, including edaphic, climatic, host genetic, and spatial factors. The detailed methods for variation partitioning are described in Matsuoka *et al.* (2016a). First, we constructed edaphic, climatic, and host genetic models by applying the forward selection procedure (999 permutations with an alpha criterion = 0.05) of Blanchet *et al.* (2008). The full models were as follows: edaphic model [pH + C/N ratio + WC]; climatic model [MAT + MAP + t2w + p2w]; and host genetic model [three variables of cluster probability]. Then, we constructed the models using spatial variables extracted based on principal components of neighbor matrices (PCNM, Borcard *et al.*, 2004). The PCNM analysis produced a set of orthogonal variables derived from the geographical coordinates of the sampling locations. We used the PCNM vectors that best accounted for autocorrelation and then conducted forward selection (999 permutations with an alpha criterion = 0.05, full model contained six PCNM variables). Based on these four models, we performed variation partitioning by calculating adjusted *R*^2^ values for each fraction (Peres-Neto *et al*., 2016).

To characterize the scale of spatial clustering in the ECM OTUs compositions, Mantel correlogram analysis was performed (‘mantel.correlog’ command in the vegan package). A Mantel correlogram draws a graph in which Mantel correlation value *rM* is plotted as a function of the spatial distance classes. A positive (and significant) *rM* indicates that for the given distance class, the multivariate dissimilarity among samples is lower than expected by chance (Borcard & Legendre, 2012). Raup-Crick dissimilarity matrix of ECM OTUs community was used for the analysis. P-values were generated by permutational Mantel test (999 permutations) and adjusted using the Benjamini-Hochberg procedure (Benjamini & Hochberg, 1995) to correct the significance level for multiple comparisons.

## Results

### Taxonomic assignment

In total, the 129,725 filtered pyrosequencing reads from 233 samples were grouped into 611 fungal OTUs with 97% sequence similarity (Supporting Information Table S1). Among them, 365 OTUs (92,651 reads) belonged to ECM fungal taxa, with 351 OTUs being Basidiomycota and 14 OTUs being Ascomycota. Each site yielded 36– 64 OTUs (an average of 48 OTUs, Supporting Information Table S4). At the family level, 365 ECM OTUs belonged to 19 families, and the common families were Russulaceae (134 OTUs, 36.7% of the total numbers of ECM fungal OTUs), Thelephoraceae (94 OTUs, 25.8%), and Cortinariaceae (45 OTUs, 12.3%). These three families accounted for 54.0–87.0% of total richness of ECM fungal OTUs at each site (Supporting Information Fig. S3). The most common OTU was *Cenococcum geophilum* (OTU_1), accounting for 44 of 233 samples (10 of the 12 sampling sites), followed by *Lactarius* sp. (OTU_563, 33 of 233 samples and 9 of the 12 sites), *Clavulina* sp. (OTU_603, 29 of 233 samples and 8 of the 12 sites), and Thelephoraceae sp. (OTU_602, 28 of 233 samples and 10 of the 12 sites). No ECM OTUs were detected from all the 12 sites. Whereas, 153 OTUs (41.9%) of the 365 ECM OTUs were detected in only one sample. Among the remaining 212 OTUs, which were detected in more than one samples, 172 OTUs were found in less than four sampling sites (Supporting Information Fig. S4) and these OTUs include those detected only in geographically proximal sites. For example, *Tomentella* sp. (OTU_81) and *Russula* sp. (OTU_591) were found only from the South-West islands (Site 1, 2, and 3, Supporting Information Table S1).

### OTU richness of ECM fungi

The OTU richness significantly increased with latitude (Pearson’s *r* = 0.2078, P = 0.0019, n = 233) and longitude (Pearson’s *r* = 0.1991, P = 0.0030, n = 233). The model selection of the GLMs by BIC concluded that the model with host allelic richness + p2w + MAP + MAT was the best model accounting for the variation in OTU richness (Supporting Information Table S5). The best model showed that the OTU richness increased with host allelic richness and with MAP and decreased with MAT and with p2w. Among the four environmental variables, MAT had the strongest relation with the OTU richness (Deviance = 40.29, pseudo-R^2^ = 0.1687, Table 2).

**Table 2.** The best generalized linear model (GLM) for the effects of environmental variables on rarefied ECM fungal OTU richness based on Bayesian information criterion (BIC).

|  | df | Deviance | coefficient | Partial<br>pseudo-R<sup>2</sup> |
| --- | --- | --- | --- | --- |
| Host allelic richness | 1 | 2.65 | 0.5554 | 0.0121 |
| p2w | 1 | 9.94 | -0.0015 | 0.0446 |
| MAP | 1 | 7.27 | 0.0003 | 0.0328 |
| MAT | 1 | 40.29 | -0.1264 | 0.1687 |
| Residual | 229 | 237.53 |  |  |

MAT mean annual temperature, MAP mean annual precipitation, p2w cumulative amount of precipitation during 2 weeks before each sampling date.

### Community structures of ECM OTUs

The Mantel test and NMDS ordination revealed the biogeographic changes in the ECM fungal compositions. The dissimilarity of ECM fungal community among samples was significantly and positively correlated with distances calculated from latitudes (Mantel’s *r* = 0.1054, P = 0.0001) and longitudes of sites (Mantel’s *r* = 0.1397, P = 0.0001). The NMDS plot showed the geospatial change of ECM OTU composition among sites (Fig. 2, stress value = 0.173). The ordination was significantly correlated with latitude and longitude (‘envfit’ function; latitude, R^2^ = 0.636, P = 0.010; longitude, R^2^ = 0.715, P = 0.006).

**Fig. 2.**
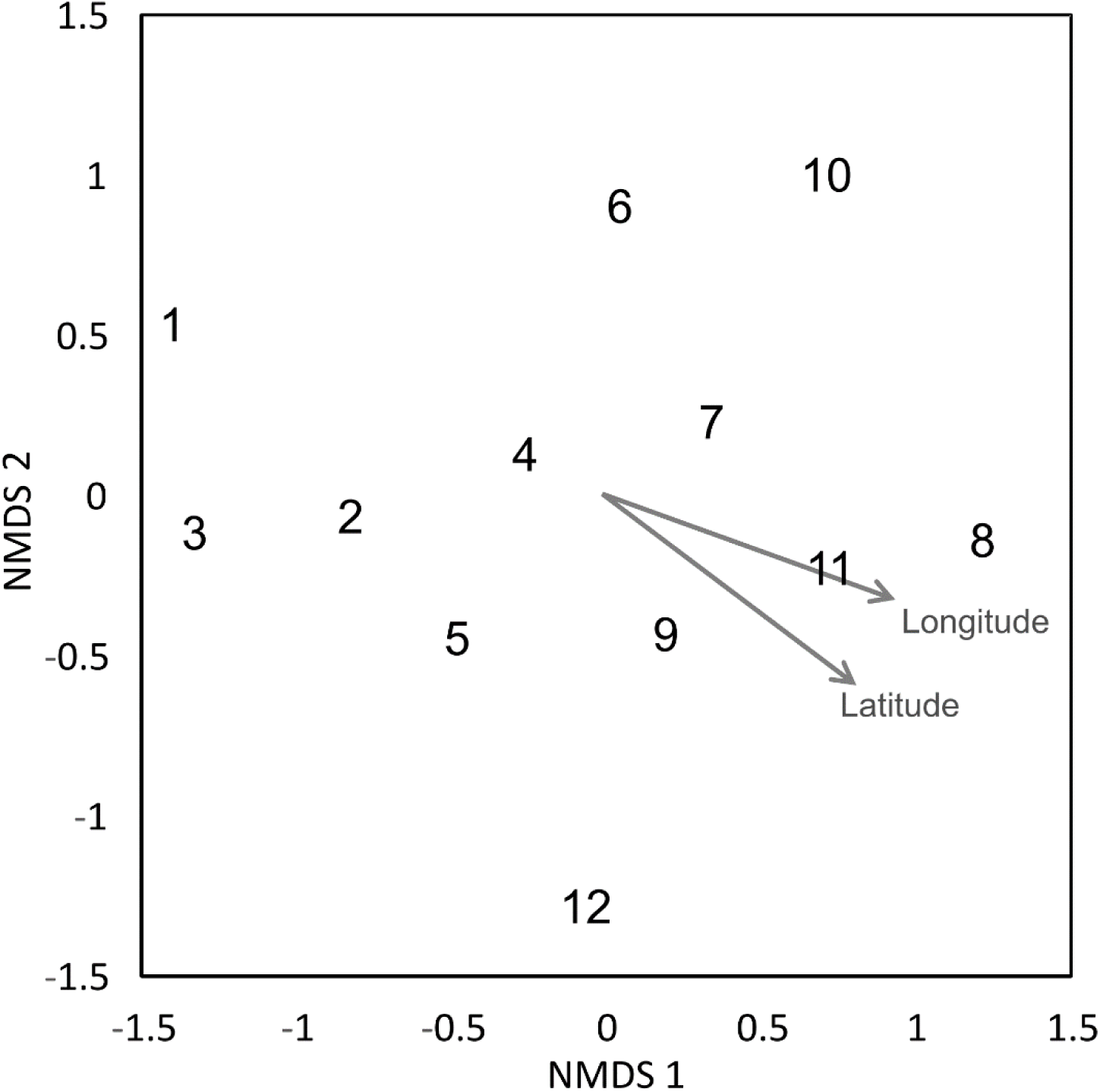
Community dissimilarity among the sites as revealed by nonmetric multidimensional scaling (NMDS) ordination (stress value = 0.173). Numbers are consistent with site numbers listed in Table 1.

In the variation partitioning, pH, C/N ratio, and WC of the FH materials were selected as edaphic factors, MAT, MAP, t2w, and p2w were selected as climatic factors, three STRUCTURE variables were selected as host genetic factors, and four PCNM vectors (vector 4, 5, 1, and 2) were selected as spatial factors. The percentages explained by the edaphic, climatic, host genetic, and spatial fractions were 6.8%, 14.6%, 10.6%, and 18.5%, respectively. Pure fractions of soil, climate, host, and space were 2.2%, 7.9%, 3.8%, and 11.2%, respectively (Fig. 3). Among the shared fractions, spatially structured host genetic fraction (i.e., shared fraction between space and host, 4.0%) and climatically structured host genetic fraction (i.e., shared fraction between climate and host, 3.7%) were the two largest fractions (Fig. 3). In total, 36.2% of the community variation was explained and the remaining 63.8% were unexplained.

**Fig. 3.**
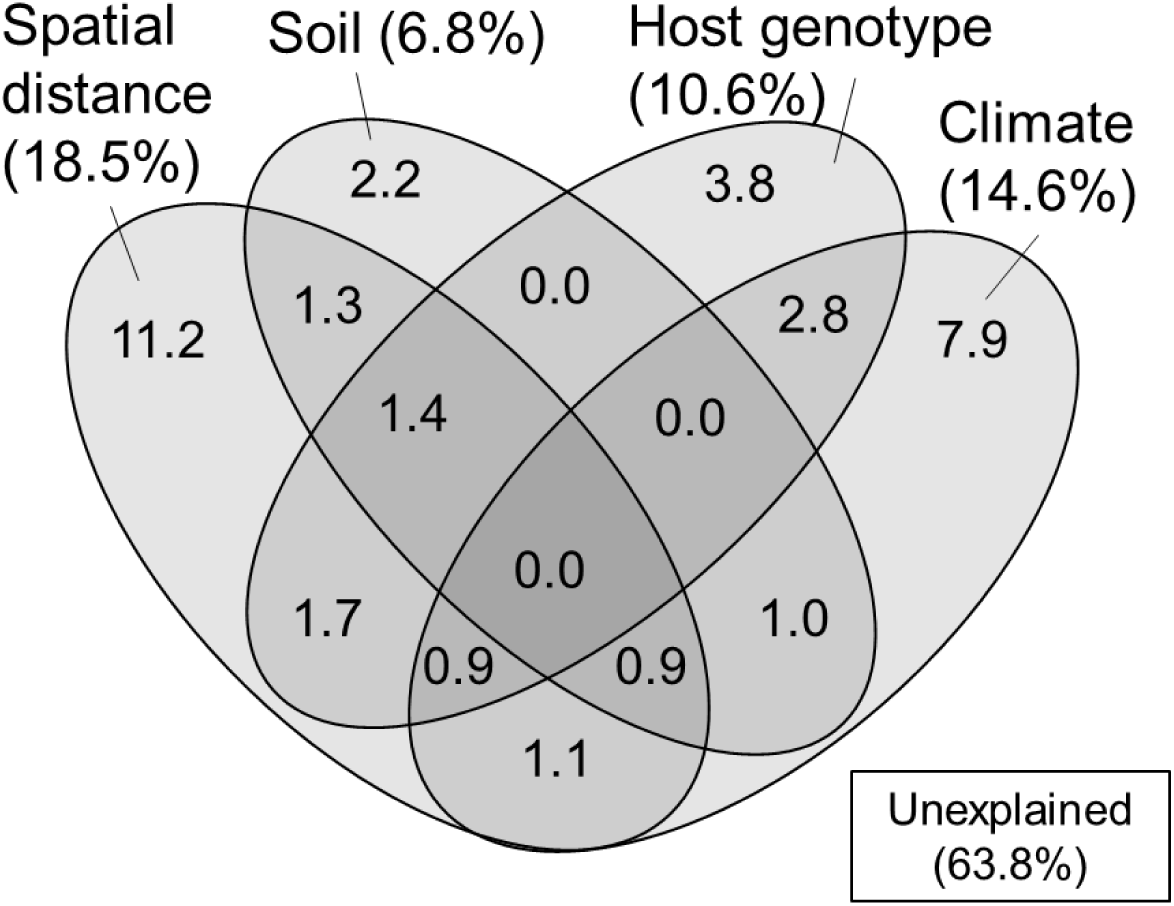
Venn diagram showing pure and shared effects of soil, climate, host genotype, and spatial distance on the ECM fungal community composition as derived from variation partitioning analysis. Numbers indicate the proportions of explained variation.

In the Mantel correlogram, in which data were lumped into nine distance classes for this analysis, the first two distance classes exhibited significant spatial autocorrelation of the ECM fungal community (P < 0.05, Fig. 4). The first and second distance classes represent 108 km and 325 km, respectively. No significant positive autocorrelations were found for the longer distance classes (P > 0.05).

**Fig. 4.**
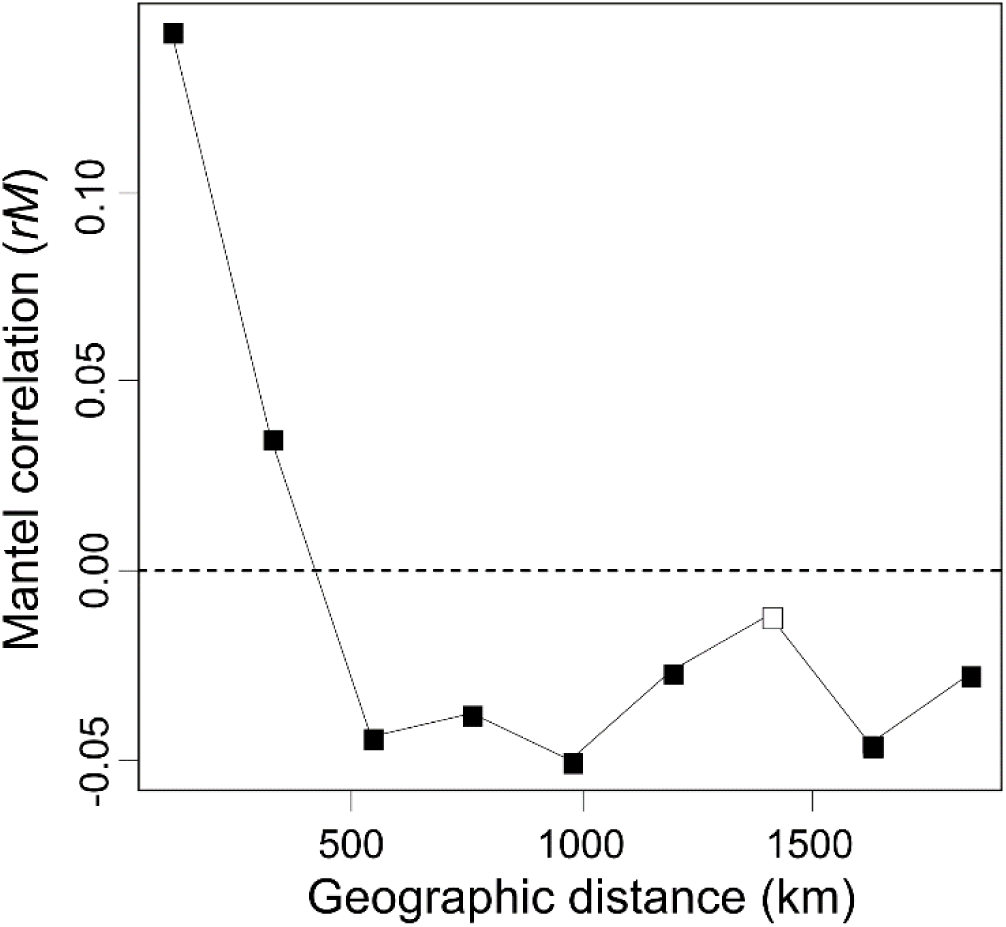
The Mantel correlogram shows the extent of spatial autocorrelation of the ectomycorrhizal fungal community composition. Filled and open boxes indicate the significant and non-significant correlation at 5% level, respectively.

## Discussion

In the present study, we successfully demonstrated the biogeographic pattern of the ECM fungal community associated with single host species (*C. sieboldii*) and detected the effects of host genetic variables on the ECM community. Our results suggest that the biogeographic pattern of ECM fungi was partly affected by the host phylogeography, which reflects the historical migration and distribution expansion of the host species. The populations of *C. sieboldii* were divided into three genetic clusters, one in the South-West Islands, another in Western, and the other in Eastern Japan (Method S1 Fig. M1-1, M1-2 and M1-3; Aoki *et al.*, 2014). These clusters indicate that, during the Last Glacial Maximum, major refugia existed in these three regions separately and *C. sieboldii* expanded its distribution from these refugia (Aoki *et al.*, 2014).

The relation between the genetic clusters of *C. sieboldii* and OTU composition of ECM fungi may be due to the past migration of ECM fungi following the expansion of their host range (i.e., co-migration) after the last glacial period. In this case, the OTU composition of ECM fungi could show spatial structure reflecting the spatial structure of genotype composition of *C. sieboldii* (i.e., geographically closer populations share similar genotype composition). Such “host mediated” spatial structure may be included in the shared fraction between host and spatial variables in variation partitioning (Fig. 3). Such co-migration of ECM fungi with their host has been reported in some previous studies, for example, current patterns of genetic relatedness in the Périgord truffle (*Tuber melanosporum*) seem to mirror patterns of post-glacial revegetation in the French Alps (Murat *et al.*, 2004). Alternatively, the preference of ECM OTUs to particular host genotypes (Gehring *et al.*, 2017; Patterson *et al.*, 2019) could also explain the relationship between host genotype and ECM composition. That is, the difference in genotype corresponds to the difference in some traits such as drought tolerance, and ECM fungi respond to these trait differences (Gehring *et al.*, 2017; Patterson *et al.*, 2019). The fraction explained only by host genetic variables in variation partitioning may include such genotype preference of ECM fungi, which is irrespective of spatial distance. Nevertheless, the *C. sieboldii* trait preference of the associated ECM fungi was evaluated only in a few studies, and the importance of preference of ECM species to particular host genotypes on ECM communities has not been fully clarified (Bubner *et al.*, 2013; Lang *et al.*, 2013).

A positive correlation was found between OTU richness and the host’s genetic diversity (i.e., allelic richness). In a group with high allelic richness, it can be inferred that a large population size has been maintained for a long time or different populations have merged (Petit *et al.*, 2003). For *C. sieboldii*, the genetic diversity was higher in sites around the three regions where refugia were speculated to be present (Supporting Information Method S1, and see above). In these sites, ECM fungal communities may have been stably maintained for a long period, providing opportunities for repeated speciation, and thus, may have yielded high species diversity. It can also be possible that the ECM OTU richness increased due to the mixing of ECM fungal communities that originated from other host populations derived from different refugia. That is, when ECM fungal communities are following the range expansion of *C. sieboldii*, at sites where different populations meet (and thus different fungal communities meet), ECM fungal diversity would become high as fungi from different communities merge. However, fungal diversity was low in low latitude region where one of the refugia was supposedly present (Supporting Information Table S4). Nevertheless, the results of GLM indicated that climatic factors should have stronger effect on ECM fungal OTU richness than host genotypic diversity.

### ECM fungal compositions and the related factors

In the present study, we successfully demonstrated that environmental filtering and dispersal limitation on fungi themselves (i.e., not via their host) can generate the biogeographic structure of the ECM community, by quantifying the effects of environmental and spatial factors on the ECM community relative to the effects of host genetic variables. Thus far, the effect of host phylogeography on biogeographic patterns of ECM communities has rarely been quantified, suggesting that patterns of ECM communities explained solely by environmental and spatial factors may also be affected by host phylogeography. Indeed, in variation partitioning, some parts of environmental and spatial fractions were shared with host genetic variables (Fig. 3). This indicates that environmental and spatial factors may partly affect the ECM community via host response to environment and dispersal, and previously demonstrated effects of these factors might be overestimated. Nevertheless, not only such shared fractions but also environmental (soil and climatic) and spatial fractions, which are not shared with the host genetic fraction, explained the difference in ECM community composition. This implies that environmental filtering and spatial processes such as dispersal limitation (Peay *et al.*, 2012; Peay & Bruns, 2014) on fungi themselves can generate the biogeographic pattern of ECM fungal community independent of the host phylogeography. Especially, the fractions explained by spatial variables alone and by climatic variable alone explained a larger proportion of the ECM composition compared to edaphic and host genetic fractions (Fig. 3). This result suggests that the biogeographic pattern of ECM community composition associated with *C. sieboldii* could be primarily structured by spatial processes reflecting historical dispersal events and by climatic filtering. Our observations are consistent with recent findings that biogeographic patterns of ECM fungal community (including ECM fungal mycelium and/or propagules in soils) are explained by spatial distance (Talbot *et al.*, 2014; Glassman *et al.*, 2015) and/or climatic variables (Bahram *et al.* 2012; Miyamoto *et al.*, 2015; van der Linde *et al.*, 2018).

It is important to note that approximately 64% of the variation remained unexplained (Fig. 3). Several factors may be invoked to explain this unexplained variation. For example, environmental factors that were not measured in the present study (e.g., soil potassium content, Tedersoo *et al.*, 2014), biotic interaction with other fungal and/or surrounding tree species (Hubert & Gehring, 2008; Bogar & Kennedy 2013; McBurney *et al.*, 2017), and stochastic processes driven by ecological drift (Hubbell, 2001) may be plausible explanations of the unexplained variation. Additionally, the spatial fraction may also result from the effects of unmeasured environmental variables that are spatially structured (Borcard & Legendre, 1994). The present study may also underestimate the explanatory power of spatial factors because our sampling sites were too sparse to detect potential spatial structures of ECM fungi in our *Castanopsis* forests. The reasons are that (i) approximately half of the OTUs in more than two samples were detected only in one sampling site, and (ii) in the result of the mantel correlogram, the distance decay of the ECM composition rapidly decreased from the first distance class (up to approximately 100 km) to the second distance class (up to approximately 300 km), which is short considering the distances between sampling sites (Fig. 1).

### OTU richness of ECM fungi

We observed that OTU richness decreased at lower latitudes. From the results of GLM, this geographic pattern of OTU richness could be attributed partly to climatic factors, especially temperature and precipitation (Table 2 and Supporting Information Table S3). Operational taxonomic unit richness decreased at sites with high MAT. This pattern is consistent with the result from a meta-analysis of global patterns of ECM communities (Tedersoo *et al.*, 2012), which showed that ECM fungal richness decreases at tropical and subtropical regions. This lower richness in tropical and subtropical region may be related to the difference in thickness of the soil organic layer. In low latitude regions, soil organic layer is thinner than that in high latitudinal regions because of active organic matter decomposition, which can make vertical habitat segregation of ECM fungi difficult (Dickie *et al.*, 2002; Tedersoo *et al.*, 2012). Indeed, it was reported from a previous study that, in Japan, the soil organic layer is thinner in sub-tropical forests than temperate forests (Osono, 2015), though in *C. sieboldii* forests, such comparison has never been conducted.

The GLM results showed that OTU richness could be related to precipitation. First, the OTU richness decreased with p2w. The potential effect of water stress (e.g., the availability of oxygen in soil decreases with increasing precipitation) on the OTU richness is discussed as a reason for the negative influence of rainfall (e.g., Tedersoo *et al.*, 2012). In our study sites, p2w had a wide range from 0 to 400 mm (Table 1), and water stress could affect OTU richness in the sites with high p2w. Whereas, MAP showed a positive relationship with OTU richness, though the explanatory value was lower than p2w. A negative relationship between OTU richness and MAP at a global-scale was reported by Tedersoo *et al.* (2012), which is contradictory to our result. The reason for this inconsistency is unclear, but the difference in the range of MAP may partly have contributed to this difference. The MAP value in our study (1,700–3,300 mm) was higher than that of the previous study (mostly around 0–2,000 mm), and this might have led to a different response of the community owing to their adaptation to high precipitation.

## Conclusions

In the present study, we demonstrated that the ECM fungal community associated with *C. sieboldii* showed a biogeographic structure that was related to the host genotypic structures, by focusing on a single host species and considering its intraspecific genetic richness and genotypes. This result indicates that the phylogeography of host species (i.e., the history of migration and environmental responses) can affect the associated ECM fungal communities and their biogeographic patterns. Moreover, with quantitative analysis, we showed that dispersal limitation and climate responses of fungi themselves, not via their host responses, could strongly affect the biogeographic pattern of the ECM fungal community. These results emphasize the importance of considering host phylogeography as well as fungal environmental responses and spatial processes (dispersal and colonization limitation) when studying the biogeographic patterns of ECM fungal community. Further studies are required to confirm whether similar patterns can be observed in other host species or climatic regions and to clarify how the relationship between intraspecific variation of host and ECM fungal community is generated.

## Acknowledgments

We thank the staff of the University of Tokyo Chiba Forests and Koichi Ito for their assistance in fieldwork; Shigeaki Hirano and Miyuki Hirata for assistance in laboratory work; Yuichiro Kanzaki for help in computer work; Ryo Kitagawa for help in statistical analyses; and Akira S Mori, Hirotoshi Sato, K. Ito for useful discussions. This study received partial financial support from the Ministry of Education, Culture, Sports, Science, and Technology of Japan (MEXT) (Grant Nos. 17K15199 and 18K05731) and the Environment Research and Technology Development Fund of Environmental Restoration and Conservation Agency (4-1602).

## Author contributions

SM designed the study. SM and TO designed and carried out the fieldwork. SM, YS, and EK performed the experiments. SM, TI, and YS analyzed the data and interpreted the results. SM, YS, HD, and TO wrote the initial draft of the manuscript. All other authors critically reviewed the manuscript.

