## Supporting Information for "Biogeographic patterns of ectomycorrhizal fungal communities associated with *Castanopsis sieboldii* across the Japanese archipelago"

The following Supporting Information is available for this article:

**Fig. S1** R Rarefaction curves of the ectomycorrhizal fungal operational taxonomic unit (ECM OTU) communities present within fermentation-humus samples as a function of sequencing read.
**Fig. S2** Proportions of major ectomycorrhizal fungal families for each study site.
**Fig. S3** Number of detection sites for ectomycorrhizal fungal OTUs which were detected in more than one sample.

**Table S2** Host genetic variables that were estimated by spatial interpolations based on the data of Aoki *et al.* (2014).
**Table S3** Correlations between geographic coordinates of the sampling sites and environmental variables used in this study.
**Table S4** Total numbers of ectomycorrhizal (ECM) fungal operational taxonomic units (OTUs) and mean values of ECM fungal OTU richness at each study site.
**Table S5** Coefficient of the factors and values of Bayesian information criterion (BIC) at each Generalized linear model (GLM).

**Methods S1** Data processing for host tree (*Castanopsis sieboldii*) genetic variables.

**Fig. S1** Rarefaction curves of the ectomycorrhizal fungal operational taxonomic unit (ECM OTU) communities present within fermentation-humus samples as a function of sequencing read.

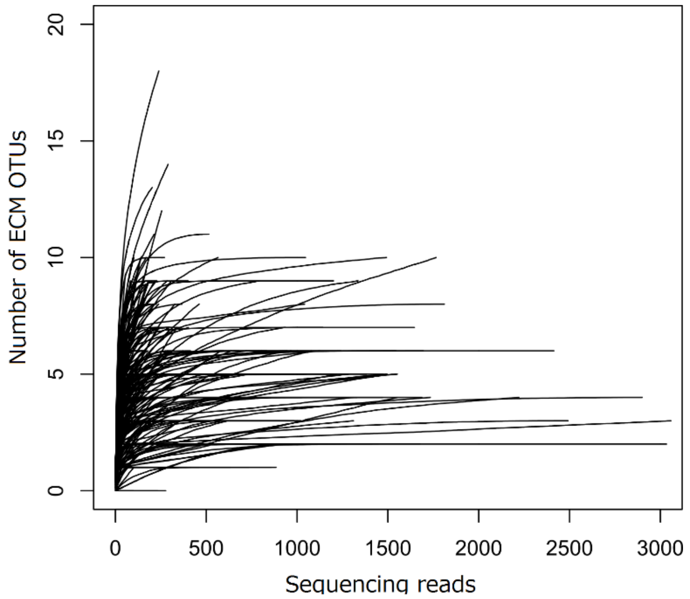

**Fig. S2** Proportions of major ectomycorrhizal fungal families for each study site.

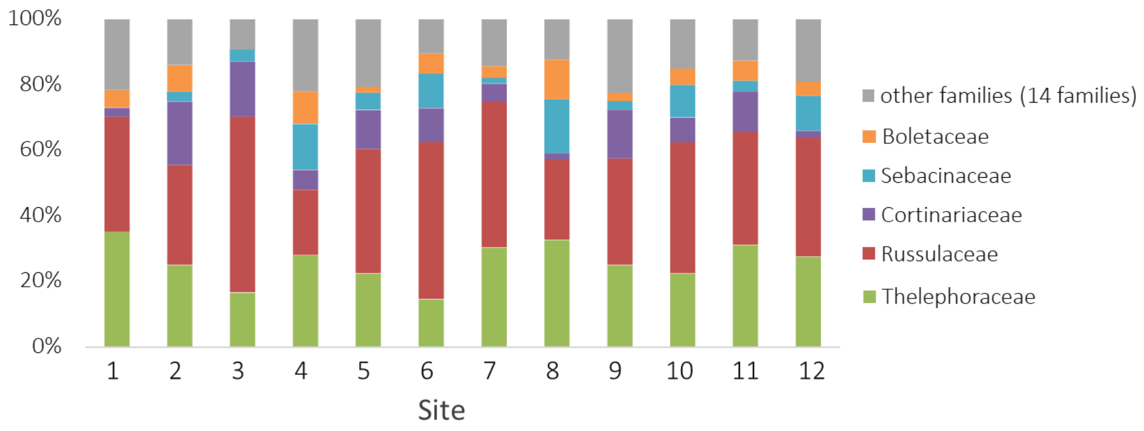

**Fig. S3** Number of detection sites for ectomycorrhizal fungal OTUs which were detected more than one sample.

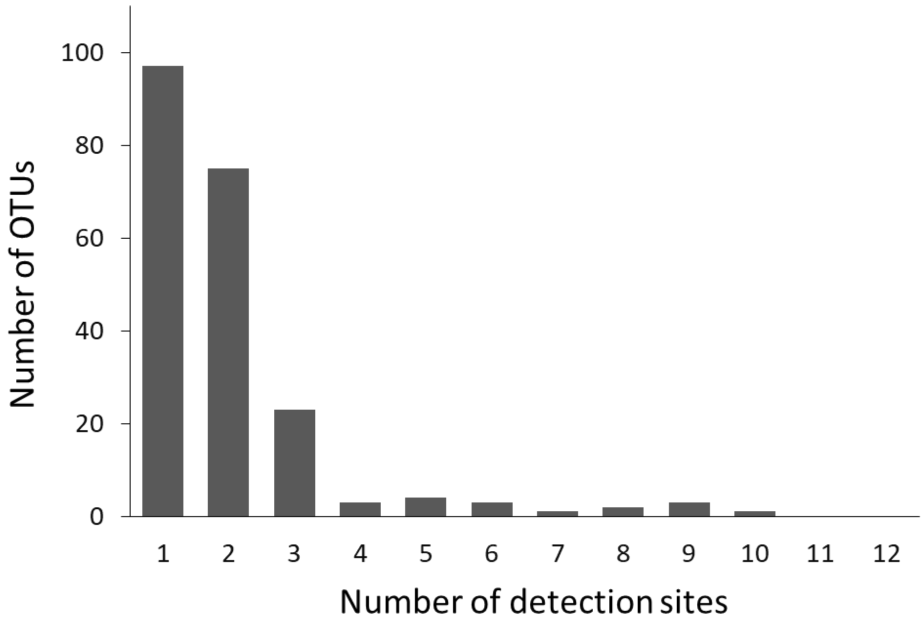

**Table S2** Host genetic variables that were estimated by spatial interpolations based on the data of Aoki et al. (2014).

| Site No. | Site name | Latitude (°N) | Longitude (°E) | Membership probability of cluster 1 | Membership probability of cluster 2 | Membership probability of cluster 3 | Allelic richness | Frequencies of rare alleles | Frequencies of private alleles |
|---|---|---|---|---|---|---|---|---|---|
| 1 | Ishigaki | 24.4178 | 124.1881 | 0.8854 | 0.0595 | 0.0570 | 5.6528 | 0.3245 | 0.0435 |
| 2 | Amami | 28.2225 | 129.3412 | 0.8478 | 0.0799 | 0.0757 | 5.6337 | 0.2491 | 0.0318 |
| 3 | Yakushima | 30.2575 | 130.5817 | 0.6787 | 0.1170 | 0.2058 | 5.5955 | 0.2245 | 0.0123 |
| 4 | Hitoyoshi | 32.1491 | 130.7803 | 0.3230 | 0.5235 | 0.1540 | 5.4728 | 0.2164 | 0.0054 |
| 5 | Saeki | 32.9594 | 131.8913 | 0.3760 | 0.4723 | 0.1536 | 5.4059 | 0.2019 | 0.0011 |
| 6 | Ashizuri | 32.7447 | 133.0002 | 0.3010 | 0.4128 | 0.2870 | 5.3539 | 0.2012 | 0.0008 |
| 7 | Kii | 33.4653 | 135.8333 | 0.1973 | 0.3011 | 0.5042 | 5.0840 | 0.1381 | 0.0017 |
| 8 | Hachijo | 33.1076 | 139.8414 | 0.1393 | 0.1164 | 0.7451 | 4.9657 | 0.1822 | 0.0158 |
| 9 | Izu | 34.6509 | 138.8526 | 0.1820 | 0.0903 | 0.7277 | 4.8719 | 0.1240 | 0.0120 |
| 10 | Chiba | 35.0934 | 139.9162 | 0.1425 | 0.0694 | 0.7882 | 5.0803 | 0.1658 | 0.0091 |
| 11 | Kurayoshi | 35.4248 | 133.8224 | 0.2038 | 0.6720 | 0.1247 | 5.1668 | 0.1143 | 0.0006 |
| 12 | Sado | 37.9661 | 138.3680 | 0.1138 | 0.7695 | 0.1171 | 4.3150 | 0.0886 | 0.0334 |

**Table S3** Correlations between geographic coordinates of the sampling sites and environmental variables used in this study.

| | | Latitude (N) | Longitude (E) |
|---|---|---|---|
| soil | pH | 0.0034 | 0.0977 |
| | Water content | 0.0310 | -0.0818 |
| | C/N ratio | **-0.2617** | **-0.3112** |
| Climate | MAT (˚C) | **-0.9570** | **-0.7191** |
| | MAP (mm) | **-0.3026** | -0.0939 |
| | t2w (˚C) | **-0.6992** | **-0.5209** |
| | p2w (mm) | **-0.2419** | **-0.4955** |
| Host | Allelic richness | **-0.8256** | **-0.8190** |
| | frequencies of rare allele | **-0.9547** | **-0.8217** |
| | frequencies of private allele | **-0.4869** | **-0.3394** |

Values are Pearson's correlation. Bold ($P < 0.05$).

MAT mean annual temperature, MAP mean annual precipitation, t2w the mean daily temperature during the 2 weeks before each sampling date, p2w cumulative amount of precipitation during 2 weeks before each sampling date

**Table S4** Total numbers of ectomycorrhizal (ECM) fungal operational taxonomic units (OTUs) and mean values of ECM fungal OTU richness at each study site. Values are mean ± S.D.

| Site No. | n[1] | Total number of ECM OTUs | Number of ECM OTUs per sample | Rarefied number of ECM OTUs per sample |
|---|---|---|---|---|
| 1 | 20 | 37 | 3.5 ± 1.6 | 2.2 ± 1.0 |
| 2 | 20 | 36 | 5.4 ± 2.7 | 3.9 ± 2.1 |
| 3 | 20 | 54 | 3.7 ± 1.5 | 3.0 ± 1.2 |
| 4 | 20 | 50 | 4.7 ± 2.2 | 3.7 ± 1.6 |
| 5 | 20 | 58 | 6.1 ± 2.6 | 5.0 ± 2.4 |
| 6 | 16 | 48 | 4.1 ± 2.1 | 3.2 ± 1.6 |
| 7 | 19 | 56 | 7.4 ± 2.5 | 4.8 ± 1.5 |
| 8 | 20 | 49 | 7.7 ± 4.0 | 6.6 ± 3.6 |
| 9 | 20 | 40 | 4.5 ± 2.0 | 3.6 ± 1.7 |
| 10 | 18 | 40 | 3.1 ± 1.3 | 2.9 ± 1.2 |
| 11 | 20 | 64 | 7.8 ± 2.3 | 6.6 ± 2.7 |
| 12 | 20 | 47 | 5.3 ± 1.9 | 3.1 ± 2.0 |

[1] the number of sample used for statistical analysis

**Table S5** Coefficient of the factors and values of Bayesian information criterion (BIC) at each Generalized linear model (GLM).

| Model No. | Intercept | allR | C/N | p2w | pH | MAP | priall | rareall | t2w | MAT | WC | df | logLik | BIC | ΔBIC |
|---|---|---|---|---|---|---|---|---|---|---|---|---|---|---|---|
| 1 | 1.041998 | 0.1040 | 0.0156 | -0.0009 | -0.2031 | 0.0005 | -6.4999 | 2.8196 | 0.0816 | -0.1696 | -0.0078 | 11 | -453.927 | 967.1833 | 20.73080 |
| 2 | 0.394621 | 0.4243 | 0.0084 | -0.0011 | -0.1867 | 0.0005 | - | 1.4116 | 0.0567 | -0.1797 | -0.0069 | 10 | -454.211 | 962.3583 | 15.90578 |
| 3 | 0.293021 | 0.4239 | - | -0.0011 | -0.2033 | 0.0005 | - | 1.1916 | 0.0660 | -0.1744 | -0.0062 | 9 | -454.396 | 957.3343 | 10.88182 |
| 4 | -0.07418 | 0.4854 | - | -0.0010 | -0.1814 | 0.0005 | - | - | 0.0601 | -0.1553 | -0.0064 | 8 | -454.545 | 952.2384 | 5.785857 |
| 5 | 1.16699 | 0.5143 | - | -0.0012 | -0.1777 | 0.0005 | - | - | - | -0.1372 | -0.0071 | 7 | -455.137 | 948.0285 | 1.576009 |
| 6 | 0.213968 | 0.5738 | - | -0.0014 | - | 0.0004 | - | - | - | -0.1335 | -0.0054 | 6 | -457.329 | 947.0199 | 0.567438 |
| 7 | 0.188086 | 0.5554 | - | -0.0015 | - | 0.0003 | - | - | - | -0.1264 | - | 5 | -459.742 | 946.4525 | 0.000000 |
| 8 | 2.071561 | - | - | -0.0010 | - | 0.0003 | - | - | - | -0.0712 | - | 4 | -466.792 | 955.1583 | 8.705755 |
| 9 | 1.959293 | - | - | - | - | 0.0003 | - | - | - | -0.0670 | - | 3 | -474.806 | 965.7936 | 19.34105 |
| 10 | 2.233342 | - | - | - | - | - | - | - | - | -0.0474 | - | 2 | -481.588 | 973.9631 | 27.51060 |
| 11 | 1.39196 | - | - | - | - | - | - | - | - | - | - | 1 | -489.817 | 985.0273 | 38.57476 |

allR allelic richness of host species, p2w cumulative amount of precipitation during 2 weeks before each sampling date, MAP mean annual precipitation, prial frequencies of private allele of host species, rareall frequencies of rare allele of host species, t2w the mean daily temperature during the 2 weeks before each sampling date, MAT mean annual temperature, WC water content of FH materials, df degree of freedom, logLik Log-likelihood.

**Methods S1** Data processing for host tree (*Castanopsis sieboldii*) genetic variables.

We estimated host genetic variables from our study sites based on the data published by Aoki *et al.* (2014). The host variables represent intra-specific differences in genotype (i.e., membership probabilities of three STRUCTURE clusters (see main text)) and diversity (i.e., allelic richness, frequencies of rare alleles, and private alleles) of *C. sieboldii* at each sampling site. The data in Aoki *et al.* (2014) could not be directly used in the present study because our 12 study sites were different from those of Aoki *et al.* (2014). Therefore, we estimated host genetic variables in our study sites by spatial interpolation in GIS software.

At first, we imported the data of the 40 study sites in Aoki *et al.* (2014) into the ArcGIS ver. 10.2 software (ESRI Products, Redlands, California). The data of allelic richness, percentages of rare alleles and private alleles in Table S1 in Aoki *et al.* (2014) were used. The STRUCTURE membership probabilities of the three clusters (cluster 1, 2, and 3 in Aoki *et al.* (2014)) were extracted from the pie charts in Figure 8 in Aoki *et al.* (2014) by the software illustrator ver. CS6 (Adobe Systems Inc., San Jose, California).

Second, we estimated the spatial distribution patterns of these genetic parameters by spatial interpolations with a kriging method in ArcGIS. The results of spatial interpolations are shown as follows (Fig. M1-1, M 1-2, M 1-3, M 1-4, M 1-5, and M 1-6). Because these six biogeographic variables (i.e., three STRUCTURE membership probabilities, allelic richness, and percentages of rare alleles and private alleles) were clearly geographically related, we used them as host biogeographic variables in the present study.

Lastly, we extracted the estimated values at our 12 study-sites using the point sampling tool in ArcGIS. The obtained data are shown in Supporting Information Table S2, and were used for the analyses in the present study.

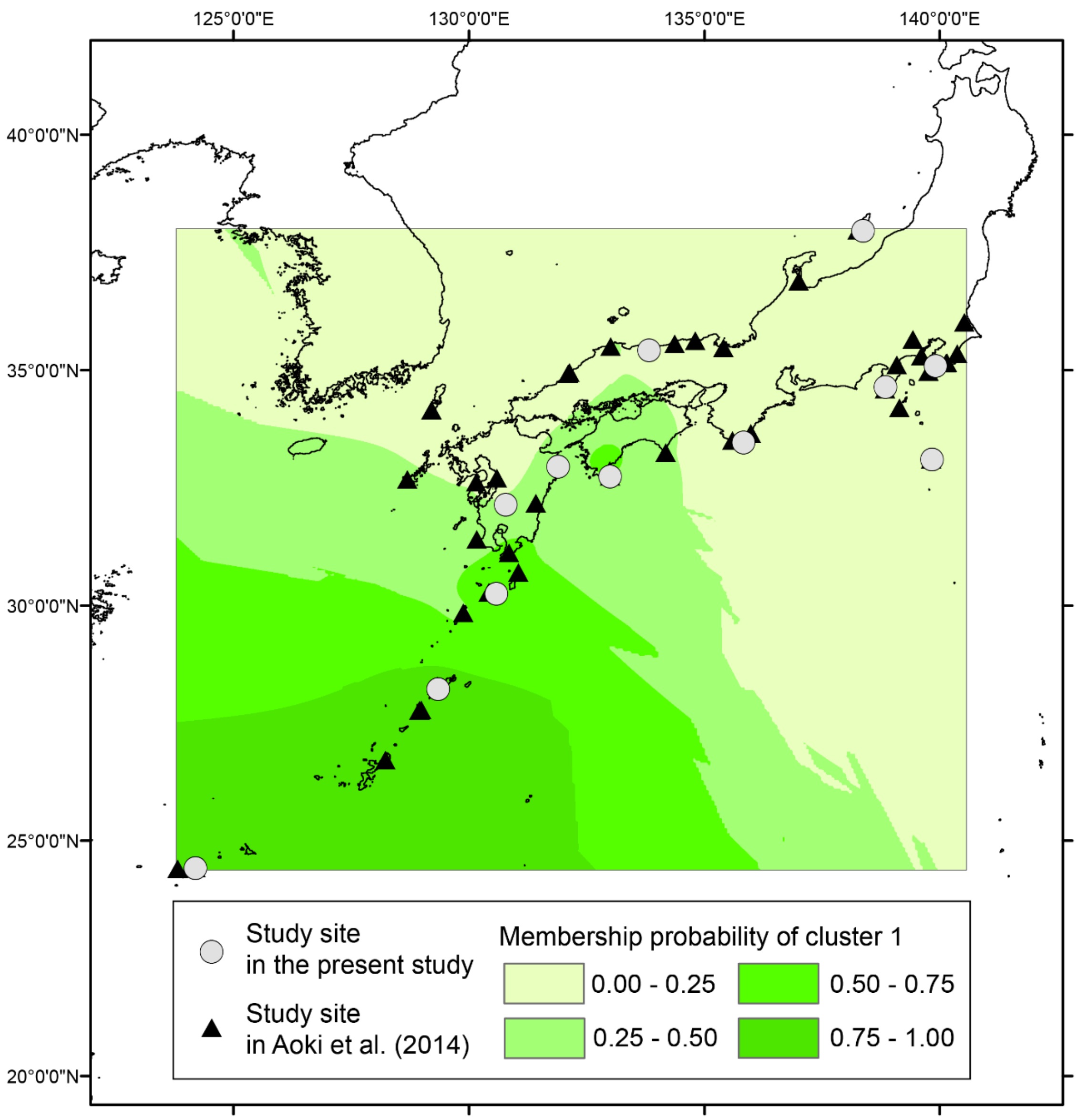


**Fig. M1-1** Result of spatial interpolation of the membership probability of STRUCTURE cluster 1.

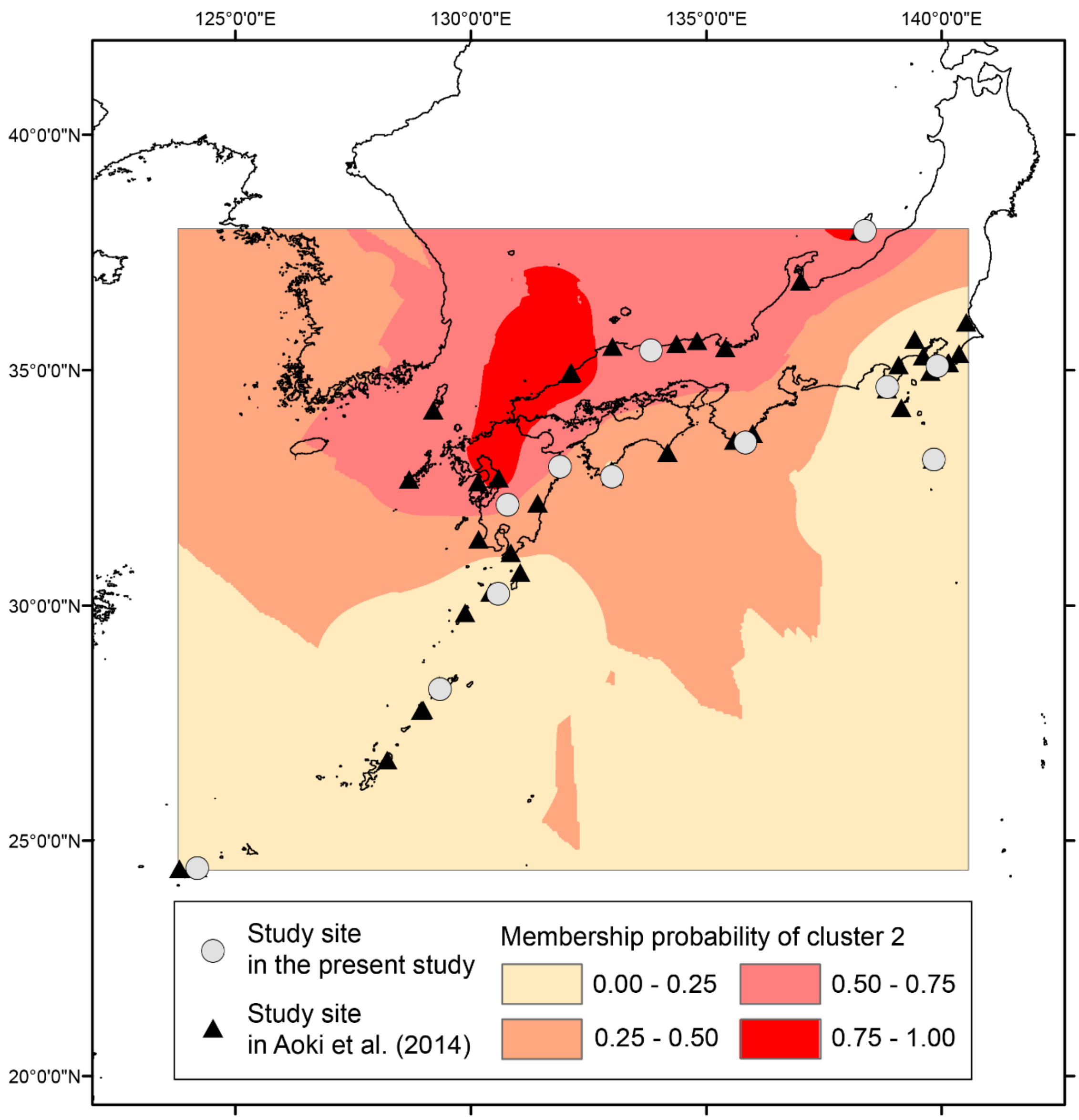


**Fig. M1-2** Result of spatial interpolation of the membership probability of STRUCTURE cluster 2.

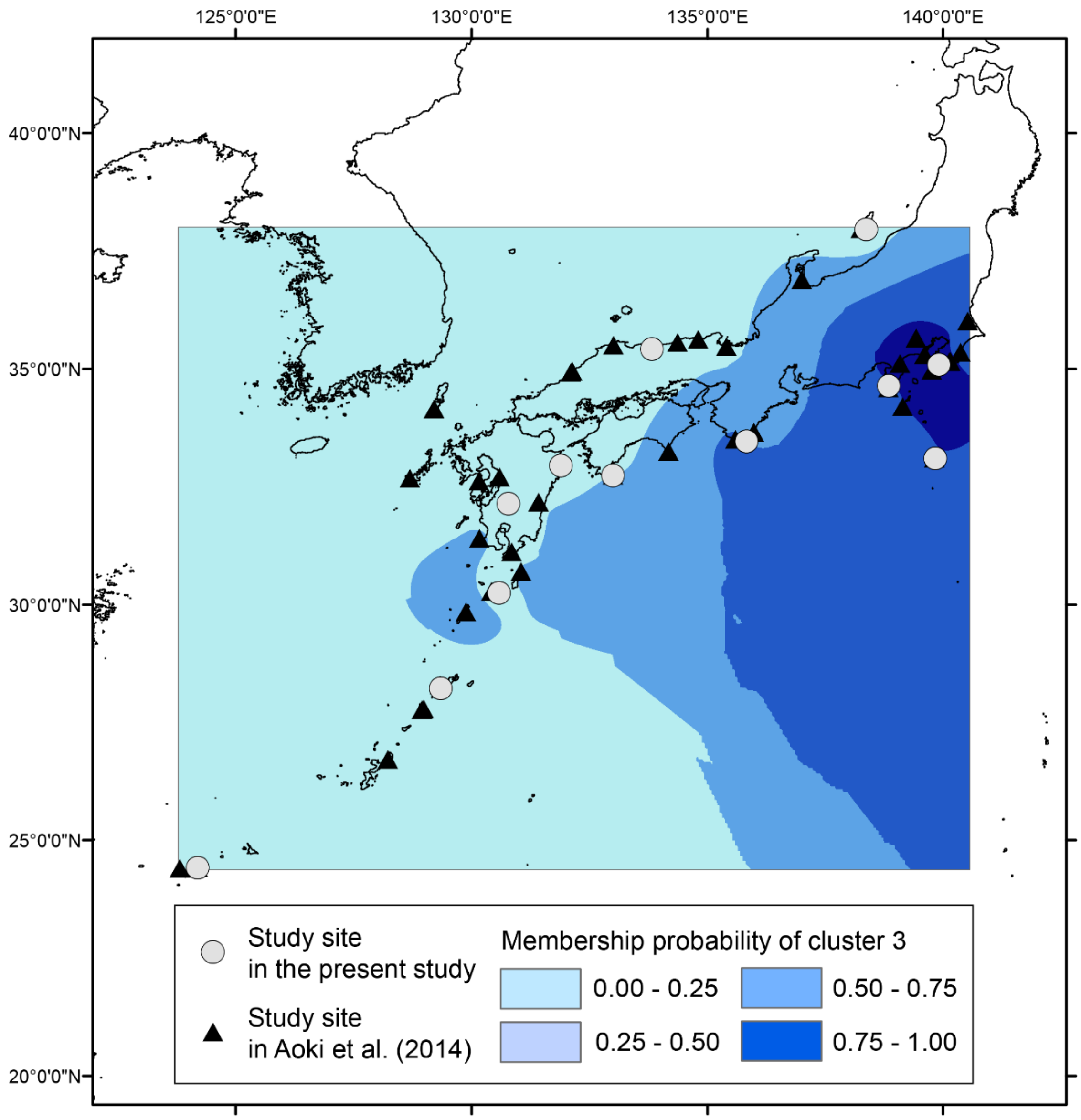


**Fig. M1-3** Result of spatial interpolation of the membership probability of STRUCTURE cluster 3.

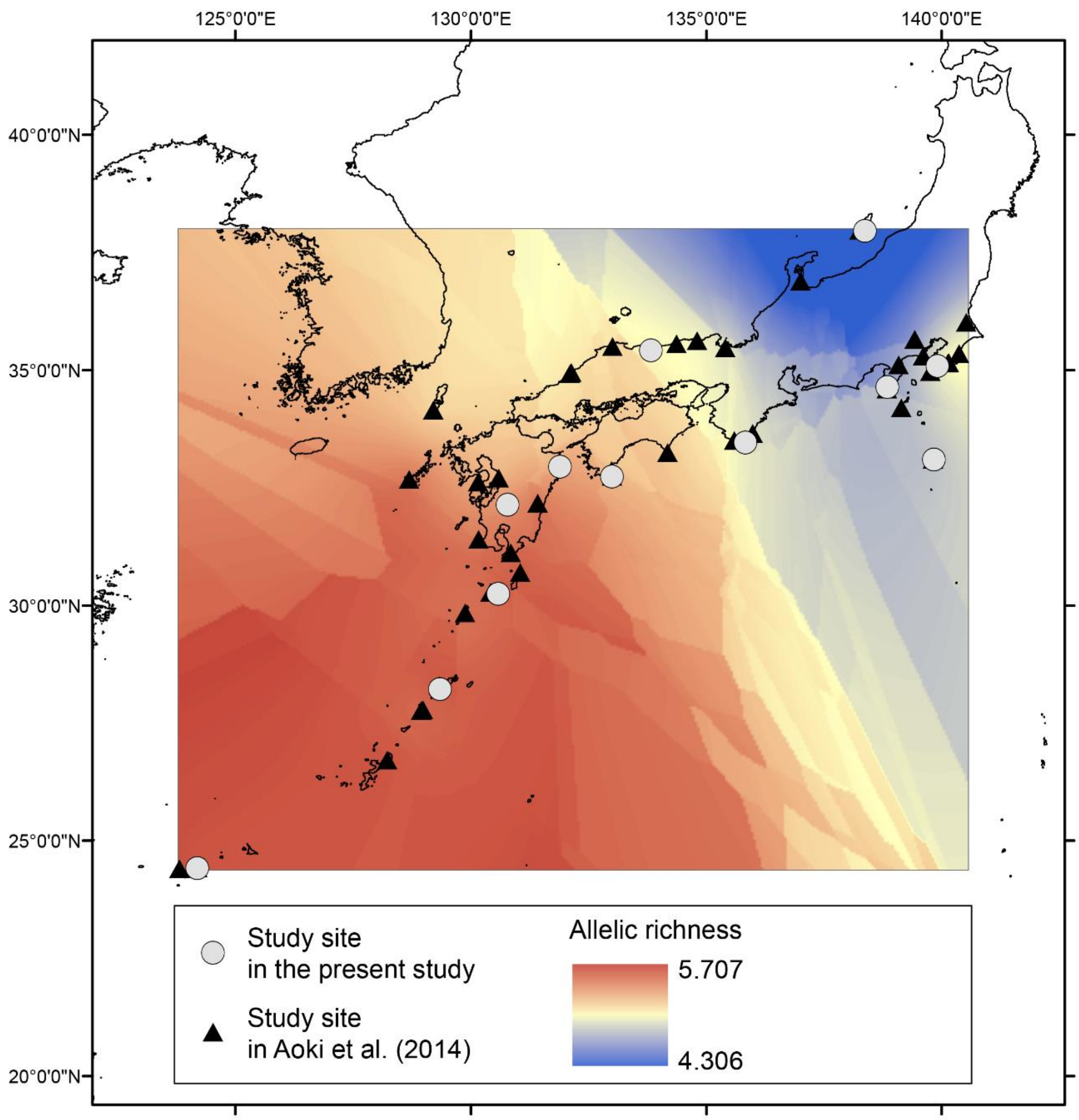


**Fig. M1-4** Result of spatial interpolation of allelic richness.

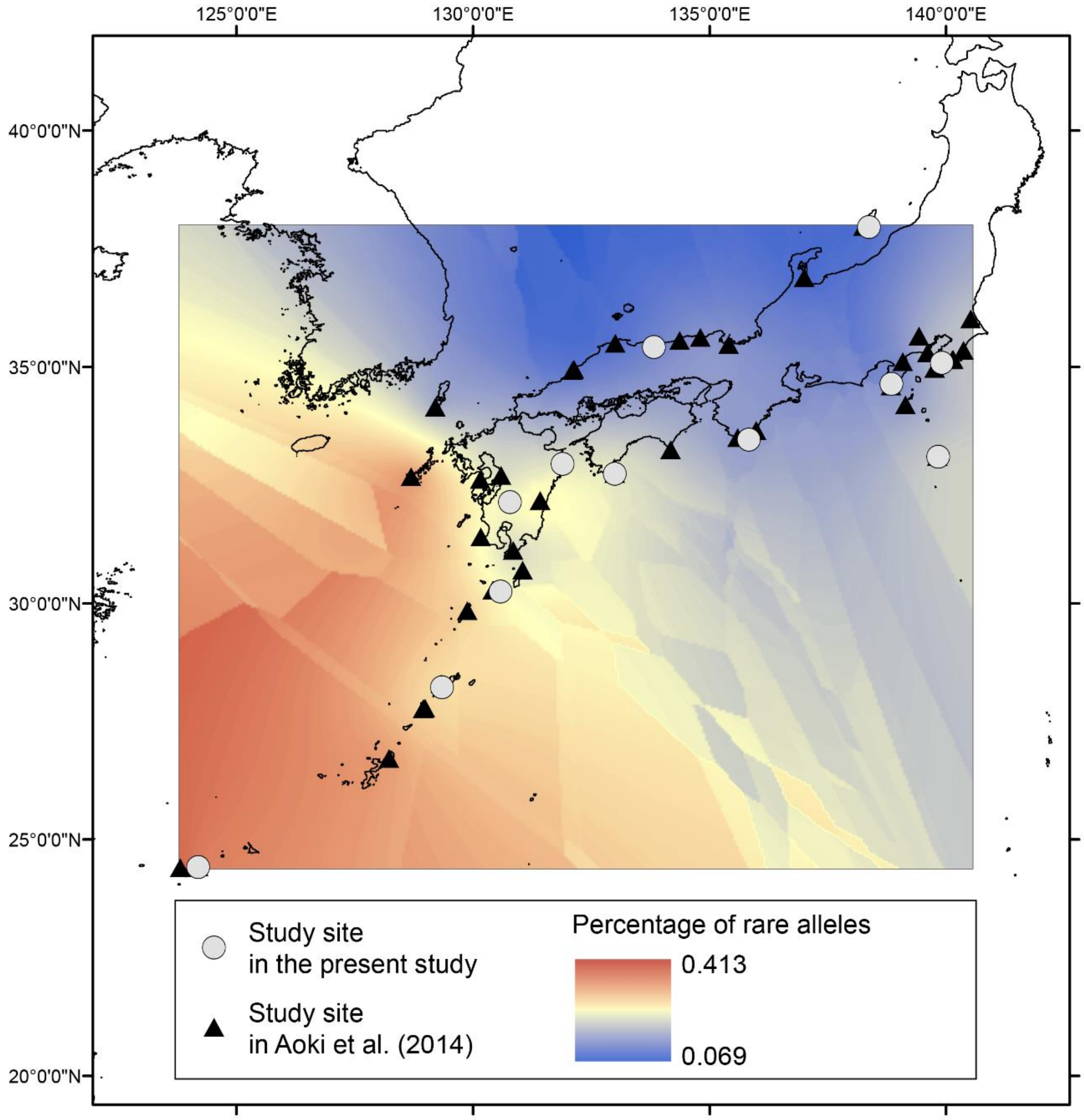


**Fig. M1-5** Result of spatial interpolation of the percentage of rare (<5%) alleles.

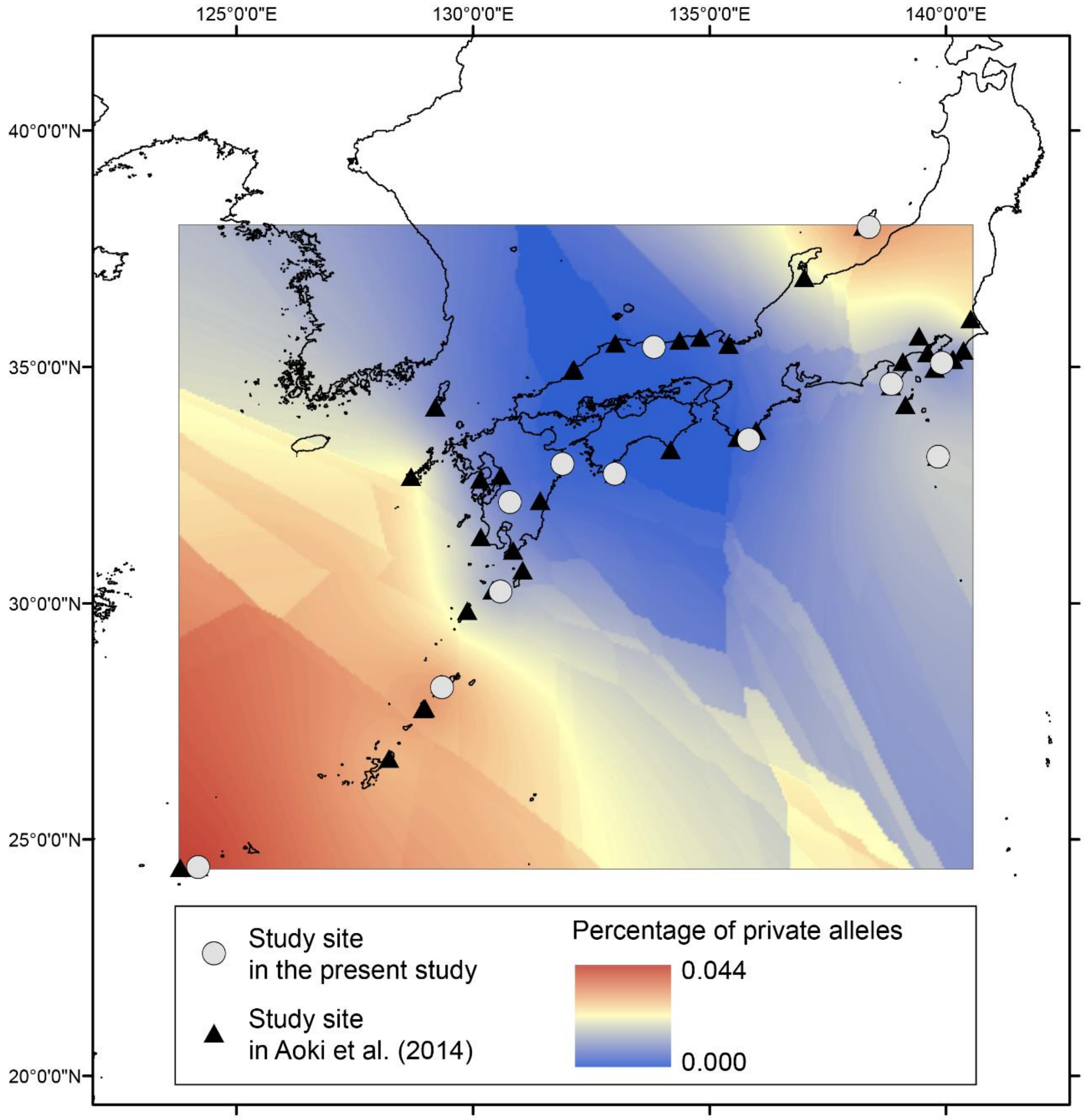


**Fig. M1-6** Result of spatial interpolation of the percentage of private alleles.
